# Plant traits that influence flower visits by birds in a montane forest

**DOI:** 10.1101/2020.08.22.262964

**Authors:** Oscar Gonzalez

## Abstract

In a bird-flowering plant network, birds select plants that present traits attractive to them. I studied plant characteristics that might predict flower visitation rate by the most common bird visitors in a bird-flowering plant network located in an elfin forest of the Andes. The nectarivorous birds which had the highest number of interactions with flowering plants in this network were the Coppery Metaltail (*Metallura theresiae*), the Great Sapphirewing (*Pterophanes cyanopterus*), and the Moustached Flowerpiercer (*Diglossa mystacalis*). I analyzed different flower traits (flower aggregation, nectar volume, nectar energy, color, orientation, and dimensions of the corolla) of the common plants that these birds visited with a principal component analysis. The plants most visited by birds were *Brachyotum lutescens* and *Tristerix longebracteatus.* While nectar traits of the plants seemed to be the best predictor for bird visitation, there was no statistical association between visitation and plant traits, except for *Metallura theresiae* in the dry season. I discuss the possible causes of resource partitioning for these nectarivorous birds.

## Introduction

Birds that feed on nectar make decisions on multiple scales to select plants and flowers; these scales could be at habitat, flowering patch, individual plant, or flower level (Sutherland and Gass 1995; Ortiz-Pulido and Vargas-Licona 2008). The visitation of each bird species may be different for the same resource (Feinsinger 1976; Davis et al. 2015). Different plant traits can attract flower visitors, such as the color of the corolla (Wilson et al. 2006), the aggregation of flowers of the plants (Fonturbel et al. 2015), the morphological matching of the feeding apparatus with the flower (Cotton 2007), flower orientation (Aizen 2003), or nectar properties.

Nectar is a primary resource for flower visitors and is a crucial determinant in interactions between animals and plants (Wiens 1989, Rathcke 1992, Cotton 2007, Janecek et al. 2012; Justino et al. 2012). The energy resource of nectar is determined by volume present and sugar concentration; animals tend to preferentially visit flowers with the most reward (Fleming et al. 2004). It is likely that nectarivorous birds - such as hummingbirds or flowerpiercers - have specific preferences for some plants depending on the nectar volume or concentration of their flowers (Hainsworth and Wolf 1976, Nicolson and Fleming 2003, Gutierrez et al. 2004, Zambon et al. 2020), and often for amino acids (Hainsworth and Wolf 1976). Although, the best sources for amino acids in hummingbirds are insects (Abrahamczyk and Kessler 2015).

The activity of flower visitors can be predicted by flower phenology (Feinsinger 1980, Stiles 1980, Murcia 1996, Rotenberry 1990, Gutierrez and Rojas 2001, Dante et al. 2013, Magilanesi et al. 2014, Gonzalez and Loiselle 2016). For example, movements of hummingbirds are known to be associated with flower blooms (Schuchmann 1999). In temperate forests, hummingbird diversity correlates with flower density, such as in Mexico (Martinez del Rio and Eguiarte 1987), Canada (Inouye et al. 1991), and the U.S. (McKinney et al. 2012). Furthermore, seasonality in the tropics is highly influential in plants and their pollinators (Cruden et al. 1983); temperature and precipitation influence local bird activity (Bourgault et al. 2010) such as foraging time and visitation rates of hummingbirds (Fonturbel et al. 2015).

In different tropical forests, several studies have shown an association of nectarivorous birds with nectar resources. Some examples of hummingbirds and their preferences by region are as follows: In Costa Rica - breeding, molt, diversity, density, and movements with blooming of their flowers (Stiles 1978, 1985, Wolf et al. 1976); in Puerto Rico - visits to flowers depend on bill size and corolla length (Kodric-Brown et al. 1984); in Bolivia - richness with flower availability (Abrahamczyk et al. 2011); and in Colombia - life cycle with nectar energy and seasonal abundance of flowers (Gutierrez et al. 2004, Cotton 2007, Toloza-Moreno et al. 2014). However studies that looked to find a relationship between hummingbirds and nectar in a landscape gave different results (Ortiz-Pulido and Rodriguez 2011). Other nectarivorous birds may select flowers by traits other than nectar such as accessibility or inflorescence size; that is the case of African sunbirds (Schmid et al. 2015). For hummingbirds, the different foraging strategies (territorial or traplining) are also important in their floral selectivity (Feinsinger 1976).

The study of nectarivorous bird communities in the neotropics provide opportunities to understand ecological interactions in different ecosystems (e.g. Rodriguez-Flores et al. 2012, Maglianesi et al. 2014) and test specific hypotheses on the drivers of these interactions, such as morphological mismatch (Vinzentin-Bugoni et al. 2014) or nectar quality and quantity (Maruyama et al. 2014). An understudied ecosystem that has an abundant nectarivorous bird community occurs in the upper montane forest of the Andes (Ramirez et al. 2007, Gonzalez 2008). In these forests, a diverse suite of hummingbirds and flowerpiercers is abundant (Gonzalez et al. 2019). However, which factors explain the patterns of plant visitation is little known in this system. Consequently, in this study the question is: Which traits of flowering plants are associated with visits by common nectarivorous birds? I hypothesize that traits associated with energy explain flower visits better than other floral traits.

## Methods

### Study Area

This research was conducted in the elfin forest in Unchog, located in the high Andes of Peru (9° 42’ 32.33” S, 76° 9’ 39.13” W; 3700 m) from 2011 to 2014. The elfin forest is considered as an ecotone between the cloud forest and the puna grassland. It has a marked seasonality of dry (May to September) and wet periods (October to March). The dry season is not devoid of rain, but it has less rain than the wet season. The temperature is cold, colder in the dry season, and the annual range varies from -1 to 15°C.

The landscape in Unchog is hilly, with small forest pockets dominated by *Weinmannia*. The non-forested area is a matrix of puna grasslands with shrubs, the most common one being *Brachyotum spp*. I sampled three sites that concentrated the most extensive groves of elfin forest (∼8 ha each), embedded in an area of 300 ha. These sites ranged from 0.6 to 1.7 Km from each other. The plant composition was very similar in the three sites (Sorensen index of similarity ranged from 0.72-0.80 among sites), so I pooled all the information on plant traits.

### Study Species

Nectarivorous birds present in the area were recorded by direct observations with binoculars. I walked inside the forest patches and forest edges, recording the birds and their visits to the flowers. I considered a visit as the moment when a bird fed on a flower or flowers of a plant, disregarding the number of flowers visited and if the visit was legitimate (pollinating) or not because this research considers the visitor’s perspective. A matrix of observed interactions, accounting for the times a bird was visiting a plant was constructed (Gonzalez and Loiselle 2016, Gonzalez et al. 2019). Birds and plants of the bird-flowering plant visitation network that were more abundant and more connected were selected to examine which plant traits predict bird interactions (Ortiz-Pulido and Vargas-Licona 2008).

The three most quantitatively important bird species that visited flowers were Coppery Metaltail (*Metallura theresiae*) – hereafter, the Metaltail; Great Sapphirewing (*Pterophanes cyanopterus*) – hereafter, the Sapphirewing, and Moustached Flowerpiercer (*Diglossa mystacalis*) – hereafter, the Flowerpiercer. The Metaltail is a small-billed, territorial hummingbird that weighs 5.07±0.09 g. and has a bill length of 12.03±0.87 mm. The Sapphirewing is a large and non-territorial hummingbird with a mass of 9.3±1.27 g. (Dunning 2007) and a bill length of 30.06±2.78 mm. The Flowerpiercer, which was the third most abundant species in terms of flower visitations, is a passerine nectar-robber with a mass of 16.2 g. (Dunning 2007) and a bill length of 10.73±1.41 mm.

### Flower Traits

I selected a subset of 13 plants that these three bird species visited to account for flower traits that might affect bird visitation. These plants had more than one interaction with birds (Gonzalez and Loiselle 2016) and were common in at least one season of the whole period of observation (Table 1). I sampled a total of 186 individual plants and an average of 14 individuals per plant species.

**Table 1.** Characteristics of plant species frequently visited by birds in the elfin forest.

| Plant species | Flower color | Flower orientation | Number of plants sampled | Mean Height (m) | SD Height | Mean Flowers in a plant | SD Flowers in a plant |
| --- | --- | --- | --- | --- | --- | --- | --- |
| *Bomarea brevis* | Red | Pendular | 6 | 0.39 | 0.20 | 2.7 | 1.7 |
| *Bomarea setacea* | Red | Pendular | 12 | 0.40 | 0.01 | 13.6 | 7.4 |
| *Brachyotum lutescens* | Green | Pendular | 18 | 0.92 | 0.60 | 36.5 | 22.1 |
| *Brachyotum naudinii* | Purple | Pendular | 10 | 0.80 | 0.60 | 30.2 | 11.6 |
| *Centropogon isabellinus* | Red | Horizontal | 6 | 0.70 | 0.01 | 19.0 | 14.3 |
| *Desfontainia spinosa* | Red | Horizontal | 18 | 1.00 | 0.01 | 17.4 | 9.3 |
| *Disterigma sp* | White | Pendular | 11 | 1.00 | 0.01 | 17.4 | 11.3 |
| *Fuchsia decussata* | Red | Pendular | 33 | 5.47 | 2.60 | 20.6 | 11.8 |
| *Gentianella fruticulosa* | Red | Pendular | 9 | 0.10 | 0.09 | 11.7 | 3.4 |
| *Passiflora cumbalensis* | Pink | Pendular | 18 | 7.13 | 2.60 | 9.3 | 3.5 |
| *Puya pseudoeryngioides* | White | Horizontal | 23 | 0.65 | 0.01 | 43.5 | 20.5 |
| *Rubus sp.* | Purple | Horizontal | 3 | 0.23 | 0.40 | 15.2 | 13.0 |
| *Tristerix longibracteatus* | Red* | Horizontal | 19 | 6.1 | 5.80 | 30.5 | 18.2 |

\* Also has yellow, but red is more predominant

To account for the availability of the flowers, I graphed the availability of the flowers in the dry season of 2014 (May, June, and July) and in the wet season of 2013 (January, February, March, and April). The resulting phenology is representative of the whole sampling period. I recorded the color of the corolla of the flowers that the birds visited (white, pink, purple, green, and red) and the orientation as horizontal or pendular (Table 1).

It is known that hummingbirds in the Andes have specific preferences for some strata in forested habitats (Gutierrez-Zamora 2008); so for each of the plants, I estimated the height where the flowers were located in relation to the ground level (Fenster et al. 2015). I also estimated flowers per individual plant as a measurement of aggregation of the resource (Dudash et al. 2011), then corolla length (Maruyama et al. 2014) and opening (Temeles et al. 2002). Nectar volume and sugar amount were also considered (Stiles and Freeman 1993; Ornelas et al. 2007). The data collected was averaged by each plant species.

### Nectar Sampling

Nectar characteristics were measured for these 13 plants (Table 2). Nectar volume in µL was measured with calibrated capillary tubes of 75 mm and the concentration in g of sugar per 100 g of solution with a refractometer that accounted for 0 to 50%, brand VEE GEE® (Kearns and Inouye 1993). Sugar constituents were not identified. There are several problems in measuring nectar, mostly due to its own variation within flowers of the same plant, time of day, and climatic conditions (Willmer 2011). The volume of nectar varied by the time of the day (McDade and Weeks 2004a) and even in flowers of the same plant (Cruden and Hermann 1983). Other studies involving measurements of nectar volume have confirmed its large variability (Baker 1975, Bolten et al. 1979, Ayala 1986, Stiles and Freeman 1993, Gutierrez and Rojas 2001, McDade and Weeks 2004a, b, Zambon et al. 2020); so the coefficient of variability for volume and concentration was considered in the analysis, as well as the largest amount of nectar (Opler 1983).

**Table 2.**
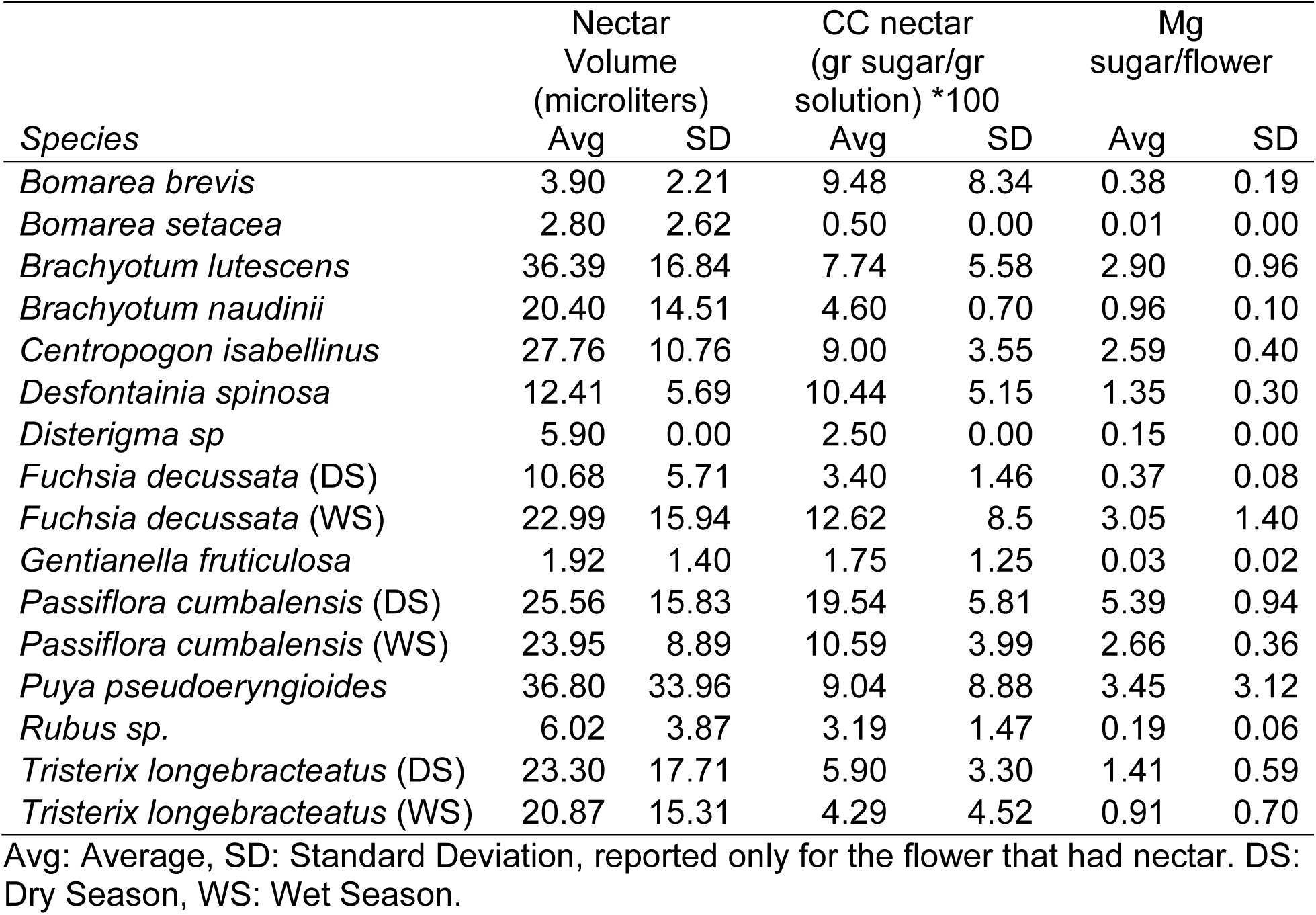
Nectar characteristics of flowers visited by birds in the elfin forest measured six hours after sunrise.

| Species | Nectar Volume (microliters) |  | CC nectar (gr sugar/gr solution) *100 |  | Mg sugar/flower |  |
| --- | --- | --- | --- | --- | --- | --- |
|  | Avg | SD | Avg | SD | Avg | SD |
| *Bomarea brevis* | 3.90 | 2.21 | 9.48 | 8.34 | 0.38 | 0.19 |
| *Bomarea setacea* | 2.80 | 2.62 | 0.50 | 0.00 | 0.01 | 0.00 |
| *Brachyotum lutescens* | 36.39 | 16.84 | 7.74 | 5.58 | 2.90 | 0.96 |
| *Brachyotum naudinii* | 20.40 | 14.51 | 4.60 | 0.70 | 0.96 | 0.10 |
| *Centropogon isabellinus* | 27.76 | 10.76 | 9.00 | 3.55 | 2.59 | 0.40 |
| *Desfontainia spinosa* | 12.41 | 5.69 | 10.44 | 5.15 | 1.35 | 0.30 |
| *Disterigma sp* | 5.90 | 0.00 | 2.50 | 0.00 | 0.15 | 0.00 |
| *Fuchsia decussata* (DS) | 10.68 | 5.71 | 3.40 | 1.46 | 0.37 | 0.08 |
| *Fuchsia decussata* (WS) | 22.99 | 15.94 | 12.62 | 8.5 | 3.05 | 1.40 |
| *Gentianella fruticulosa* | 1.92 | 1.40 | 1.75 | 1.25 | 0.03 | 0.02 |
| *Passiflora cumbalensis* (DS) | 25.56 | 15.83 | 19.54 | 5.81 | 5.39 | 0.94 |
| *Passiflora cumbalensis* (WS) | 23.95 | 8.89 | 10.59 | 3.99 | 2.66 | 0.36 |
| *Puya pseudoeryngioides* | 36.80 | 33.96 | 9.04 | 8.88 | 3.45 | 3.12 |
| *Rubus sp.* | 6.02 | 3.87 | 3.19 | 1.47 | 0.19 | 0.06 |
| *Tristerix longibracteatus* (DS) | 23.30 | 17.71 | 5.90 | 3.30 | 1.41 | 0.59 |
| *Tristerix longibracteatus* (WS) | 20.87 | 15.31 | 4.29 | 4.52 | 0.91 | 0.70 |

Avg: Average, SD: Standard Deviation, reported only for the flower that had nectar. DS: Dry Season, WS: Wet Season.

I removed nectar at different times for different flowers to check which measurement best would account for the nectar available to the plant’s potential flower visitors. I did not use the standard procedure of bagging flowers for 24 hours because there were flowers that did not produce nectar continuously, so these measurements could be misleading (Cruden and Hermann 1983; McDade and Weeks 2004a). Temperatures during the night often dropped below freezing, which causes flowers to produce less nectar. Furthermore, due to atmospheric cold fronts which are very common in this region, flower abortion is frequent; several flowers wilted or were without nectar (“rewardless”) in the early morning (59% of 929 measurements of flowers resulted in no nectar). Flowers that were covered for 6 hours since sunrise had the lowest proportion of flowers without nectar (57%). Hence, I selected this measurement as the most accurate and the best indicator for the offer of nectar to the birds. Other researchers, such as Handelman and Kohn (2014), also used nectar measurements in the morning (between 8 to 12 PM) to account for the energetic offer of the plants to hummingbirds. The standing crop (nectar mass in milligrams) for each plant species was calculated by multiplying the concentration by the volume of nectar, related by the number of hours it was covered (Cruden and Hermann 1983). Conversions were made following Dafni (1992:148).

### Analysis

I analyzed characteristics of flowers available and bird visits in wet and dry seasons separately by pooling the data across months that represented the dry season (May to September) and the wet season (October to April). I used principal component analysis (PCA) to analyze the patterns of the traits of the selected plant species (Gutierrez-Zamora 2008). This analysis identified aggregation tendencies of flower morphology (corolla length and width), distribution of flowers in the plant (flower aggregation), and flower reward to visitors (nectar volume, sugar of nectar). I used the package Factomine in R (Le et al. 2008), which helps to analyze data with multiple variables that could be numerical, ordinal, or categorical. For each of these variables, the program calculates the correlation coefficient between them and each of the values given by the plants. In this case, I set up nectar volume, sugar amount, coefficient of variance of both corolla length, corolla wide variables as numerical. The orientation of the flower (horizontal or pendular) and flower color were considered as categorical. The replicates were each one of the 13 plant species.

These plant species were ordinated based on their floral traits, such that the dispersion of the plants in the ordination reflects their separation in floral characteristics. The relative importance of the various floral characters in separating plants along the principal coordinate axes is defined by comparing the variance of the trait in the ordination with the variance of all the traits in the plot using a T-test (Le et al. 2008).

To confirm a possible association of the visitation of each species with plants of specific characteristics, I correlated the visitation data of each bird species to the plants with each of the first two axes of the PCA ordination in the dry season and in the wet season. For the comparison of bird visitations of the Sapphirewing and the Flowerpiercer with each axis, I used Spearman’s rank-order correlation due to the non-normality of the data. The statistics was done with the package Stats in R. For the visitations of the Metaltail, I performed a generalized linear model (GLM) with a Poisson distribution (Zuur et al. 2009), using the axis of the PCA as independent variables.

## Results

### Principal Components of Flower Traits

The two principal axes of the ordination accounted for 66% of all variation (Figure 1 and Sup. Table 1). Figure 1 shows only the dry season due to the ordination of plant traits was almost identical for both seasons. Plants that had greater energy (mg. sugar per flower, nectar per flower, maximum nectar and maximum concentration) and larger number of flowers per plant tended to cluster with higher scores on the first principal component axis (e.g. *Tristerix longebracteatus, Centropogon isabellinus, Brachyotum lutescens*). Plants located in higher vegetation strata - with larger corolla and wider corolla opening (this last trait becoming important only in the wet season) and few flowers per plant - tended to have higher scores along the second PCA axis (*Fuchsia decussata, Desfontainia spinosa, Passiflora cumbalensis*) (Figure 2). These results were largely consistent between the wet and dry seasons, even with some turnover in plant species that flowered.

**Figure 1.**
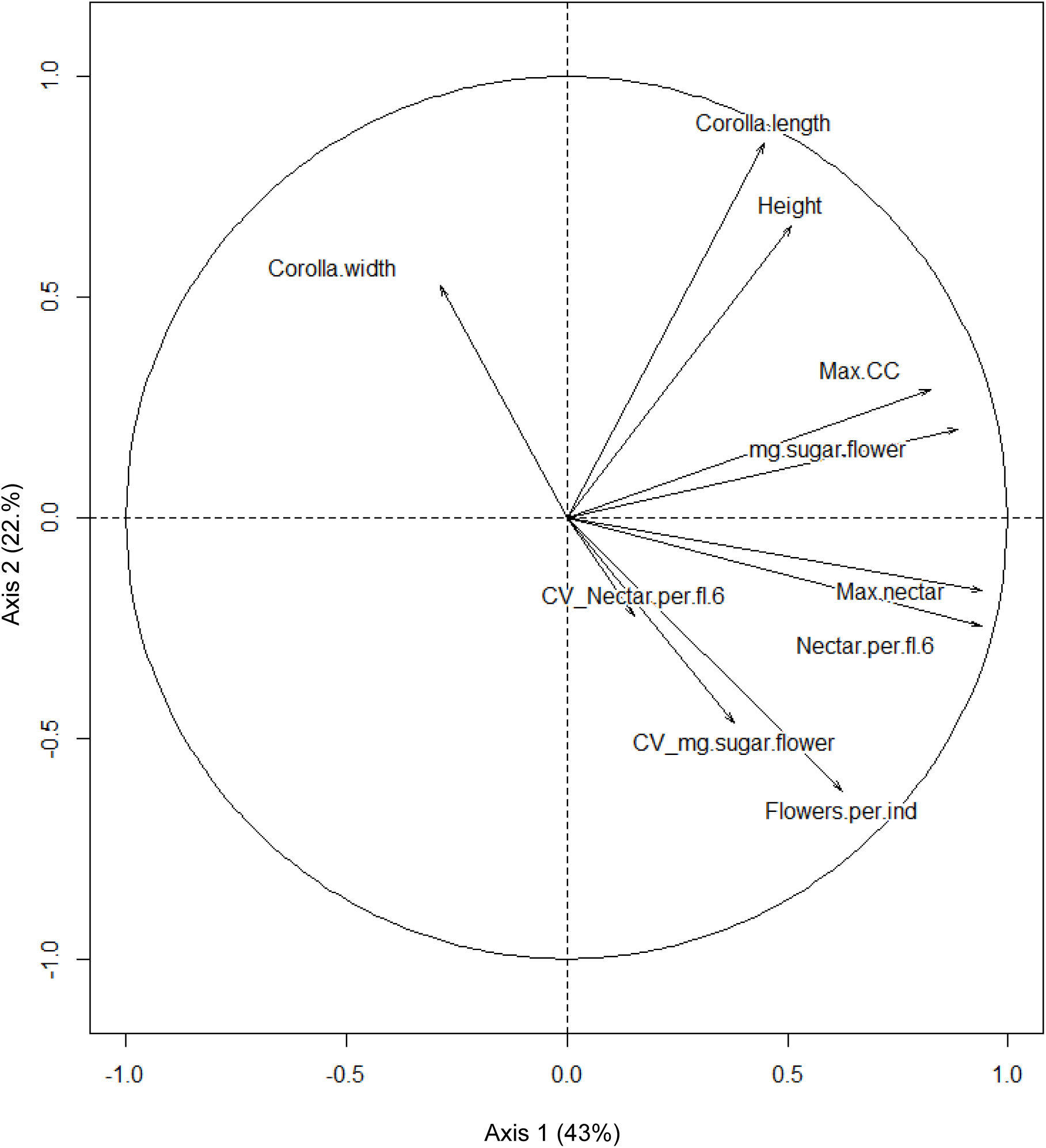
Principal component analysis of the plant traits that influence the visitation rate of the most connected bids in the bird-visitation network (dry season). Axis 1: Nectar amount (volume and mass), corolla length, height of the flower. Axis 2: Flower aggregation, nectar variability, corolla opening. Wet season was almost identical.

**Figure 2.**
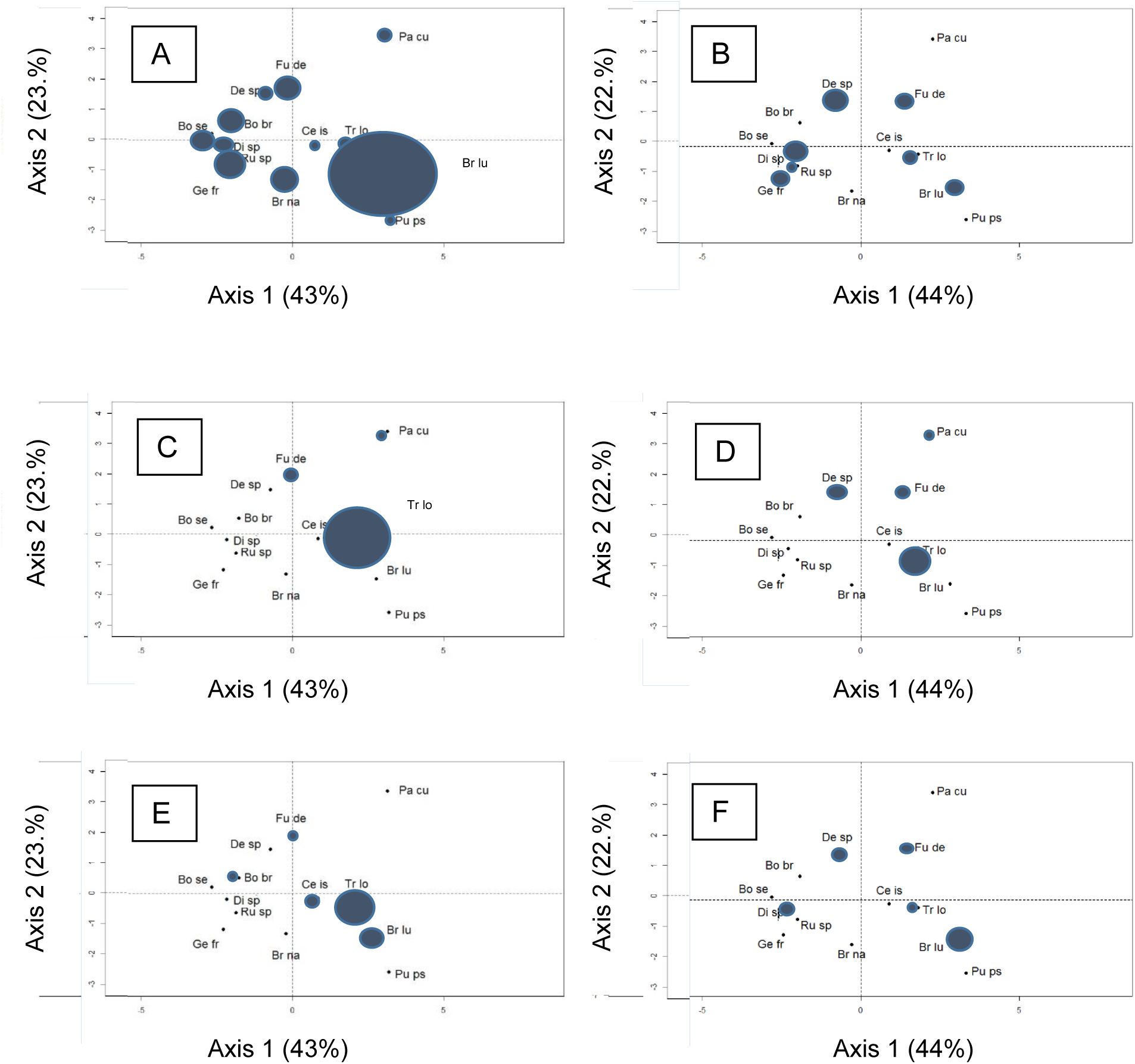
Ordination plot of the plants that influence the visitation rate of the most connected bids in the bird-visitation network. Axis 1: Nectar properties. Axis 2: Flower aggregation and morphology. The circles represent the number of visitations by birds to each plant; minimum: 1, maximum: 42. A: Metaltail, dry season. B: Metaltail, wet season. C: Sapphirewing, dry season. D: Sapphirewing, wet season. E: Flowerpiercer, dry season. F: Flowerpiercer, wet season. Keys: Bo br= *Bomarea brevis*, Bo se= *Bomarea setacea*, Br lu= *Brachyotum lutescens*, Br na= *Brachyotum naudinii*, Ce is= *Centropogon isabellinus*, De sp= *Desfontainia spinosa*, Di sp= *Disterigma sp*., Fu de= *Fuchsia decussata*, Ge fr= *Gentianella fruticulosa*, Pa cu= *Passiflora cumbalensis*, Pu ps= *Puya pseudoeryngioides*, Ru sp= *Rubus sp*., Tr lo= *Tristerix longebracteatus*.

The Metaltail in the dry season had almost half of its visitations to the shrub *Brachyotum lutescens* (Table 3), which has relatively moderate number of flowers per plant and high variability in nectar volume and sugar. The rest of their flower visits were dispersed and included plants with relatively low nectar rewards and plants that occurred in lower vegetation strata (Figure 2A). In the wet season, the Metaltail visited a greater diversity of plants as measured by their floral traits, demonstrated by their overlap in all quadrants of the ordination (Figure 2B). The Sapphirewing tended to visit plants with higher energy rewards, large corolla, and higher vegetation strata such as the mistletoe *Tristerix longebracteatus*, with 92% of all visits in the dry season (Figure 2C). Similarly, visits during the wet season were also concentrated on plants with these same characteristics. As in the dry season, the mistletoe dominated among plant visits (75%) (Figure 2D). The Flowerpiercer tended to also visit plants primarily with high nectar reward and a high number of flowers per individual such as the previous mistletoe (58% of visits) and *Brachyotum lutescens* (25% of visits) in the dry season (Figure 2E). Although *Brachyotum lutescens* accounted for 50% of the visits in the wet season (Figure 2F), like the Metaltail, flowerpiercers visited a diversity of plants across the entire ordination space.

**Table 3.** Total visitation recorded by the most connected species in the bird-flowering plant visitation network.

| Plant species/Bird visitor<br>Season | Metaltail |  | Sapphirewing |  | Flowerpiercer |  |
| --- | --- | --- | --- | --- | --- | --- |
|  | Dry | Wet | Dry | Wet | Dry | Wet |
| *Bomarea brevis* | 7 | 0 | 0 | 0 | 1 | 0 |
| *Bomarea setacea* | 5 | 0 | 0 | 0 | 0 | 0 |
| *Brachyotum lutescens* | 42 | 3 | 0 | 0 | 6 | 4 |
| *Brachyotum naudinii* | 7 | 0 | 0 | 0 | 0 | 0 |
| *Centropogon isabellinus* | 1 | 0 | 0 | 0 | 2 | 0 |
| *Desfontainia spinosa* | 3 | 4 | 0 | 2 | 0 | 1 |
| *Disterigma sp* | 4 | 1 | 0 | 0 | 0 | 1 |
| *Fuchsia decussata* | 7 | 3 | 1 | 2 | 1 | 1 |
| *Gentianella fruticulosa* | 0 | 3 | 0 | 0 | 0 | 0 |
| *Passiflora cumbalensis* | 3 | 0 | 1 | 1 | 0 | 0 |
| *Puya pseudoeryngioides* | 1 | 0 | 0 | 0 | 0 | 0 |
| *Rubus sp.* | 6 | 4 | 0 | 0 | 0 | 0 |
| *Tristerix longebracteatus* | 2 | 2 | 23 | 15 | 14 | 1 |

I found that bird visits for the Sapphirewing and the Flowerpiercer could not be explained by floral traits (Table 4). All correlations between these two birds and PCA scores for plant traits were non-significant, except for a positive correlation for the Sapphirewing along the second axis in the wet season (Table 4). However, I found a significant positive association along axis 1 and a significant negative association along axis 2 for visits by Metaltails in the dry season (Axis 1: 0.19, p<0.001; Axis 2: -0.17, p< 0.05, df=12) but not in the wet season. (Axis 1: -0.04, p=0.65; Axis 2: 0.01, p=0.92, df=12) Therefore, in the dry season, Metaltails appeared to frequently visit plants with higher energy rewards (axis 1) and plants with smaller corollas located lower in the vegetation.

**Table 4.**
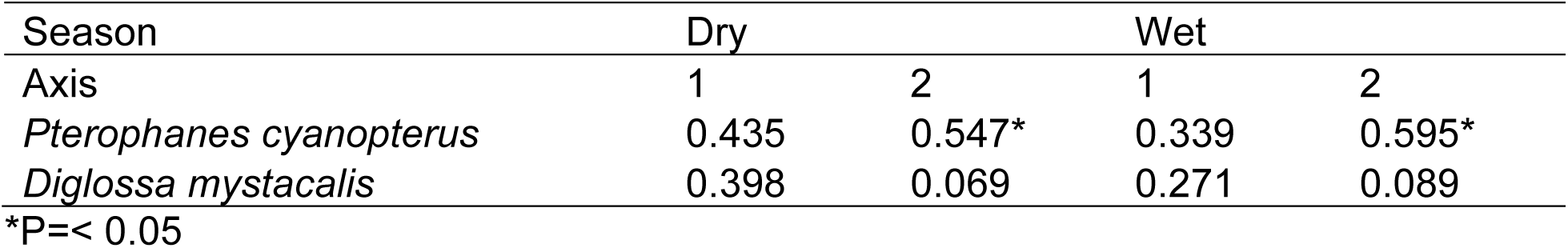
Spearman correlation coefficients for bird visitation against nectar traits (axis 1) and corolla morphology (axis 2) of the principal component analysis for flowering plants visited by birds in the elfin forest.

| Season | Dry |  | Wet |  |
| --- | --- | --- | --- | --- |
|  | 1 | 2 | 1 | 2 |
| *Pterophanes cyanopterus* | 0.435 | 0.547* | 0.339 | 0.595* |
| *Diglossa mystacalis* | 0.398 | 0.069 | 0.271 | 0.089 |

\*P=< 0.05

### Plant Trait Variability

The abundance of flowers and number of species flowering varied across seasons (Figure 3). Plants that flowered across seasons included *Bomarea brevis, Brachyotum lutescens, Tristerix longebracteatus, Fuchsia decussata* and *Rubus sp.*, while *Centropogon isabellinus* and *Bomarea setacea* produced flowers for only limited periods. *Puya* was spatially patchy and flowered only over a short period of time in the wet season. Some species, such as *Brachyotum naudinii* and *Bomarea setacea*, also flowered only in the wet season.

**Figure 3.**
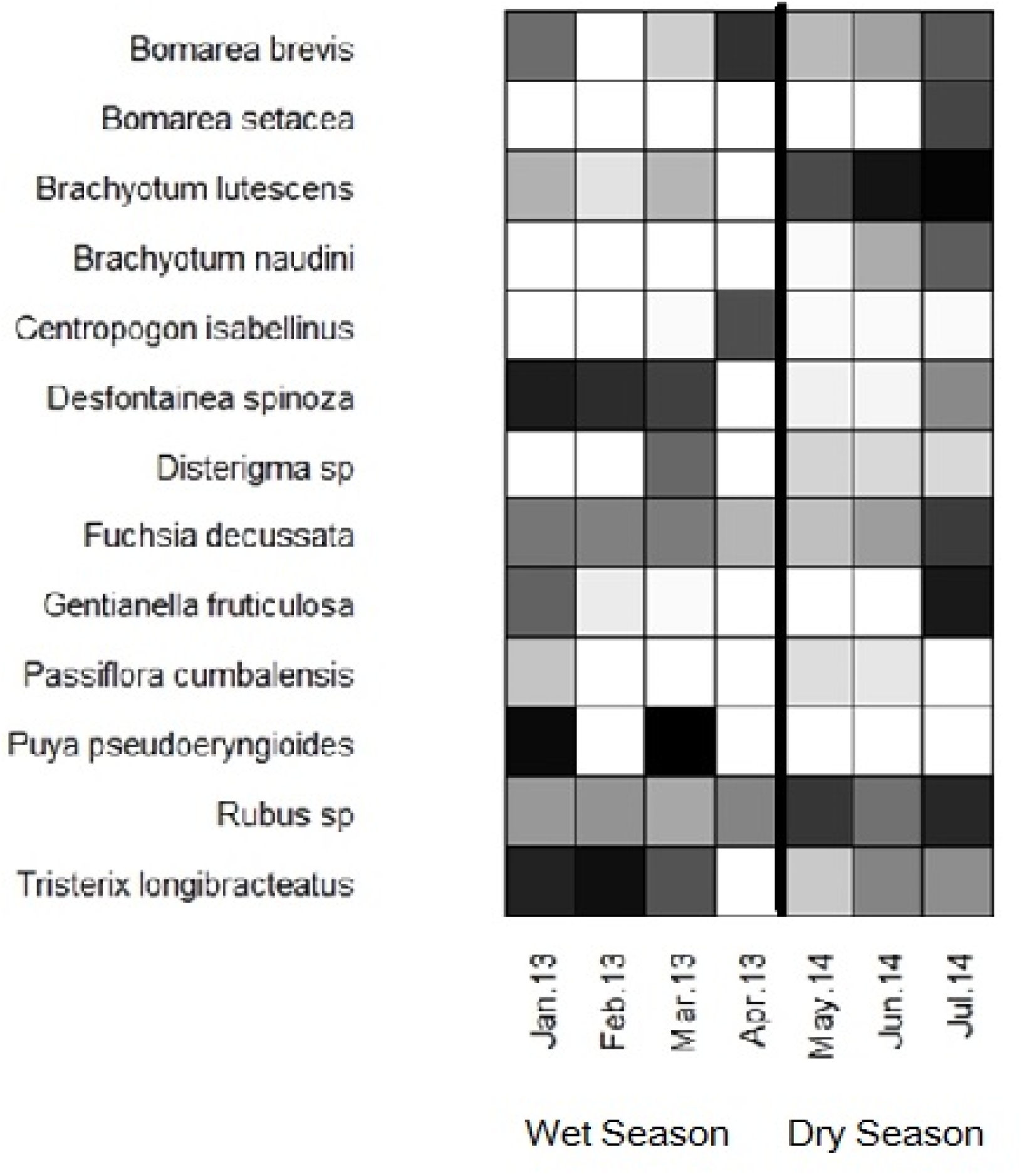
Flowering phenologies of ornithophilus plants most visited by birds in the elfin forest (2013-2014). The darker the square, the higher the number of flowers per hectare.

The factors of flower aggregation, corolla color, and flower orientation were not independent; they were linked to specific species of plants that the birds visited, so there is no way to account for floral selectivity based on these factors. Flowers of *Tristerix longebracteatus*, which are red, were visited by the Sapphirewing and the Flowerpiercer, but not by the Metaltail (Table 3). The three birds visited species of plants that had many flowers per individual (*B. lutescens* and *T. longebracteatus*); but differed in the orientation of the flowers they foraged. The Metaltail visited mostly the pendular flowers of *B. lutescens*; the Sapphirewing and the Flowerpiercer frequented horizontals of *T. longebracteatus*. The Sapphirewing foraged almost exclusively in the tree canopy, the Metaltail mostly in the understory, and the flowerpiercer was between the canopy and the understory (Sup.Table 1).

As expected, the nectar volume and concentration varied considerably among the plant species selected. I reported the information for the plants that had nectar (Table 2). Several of these species had less than 50% of their flowers with nectar (Figure 4). Further, nectar volume and concentration were also found to vary between dry and wet season for *Fuchsia decussata, Passiflora cumbalensis* and *Tristerix longebracteatus*; all three of these species had long corollas. *Passiflora cumbalensis* had the highest energy available per flower (5.39±2.66 mg) and highest concentration (19.54±5.81), followed by *Puya pseudoeryngioides, Fuchsia decussata* and *Brachyotum lutescens. Brachyotum lutescens* had higher average nectar volume, followed by *Puya* (36.39±16.84 µl).

**Figure 4.**
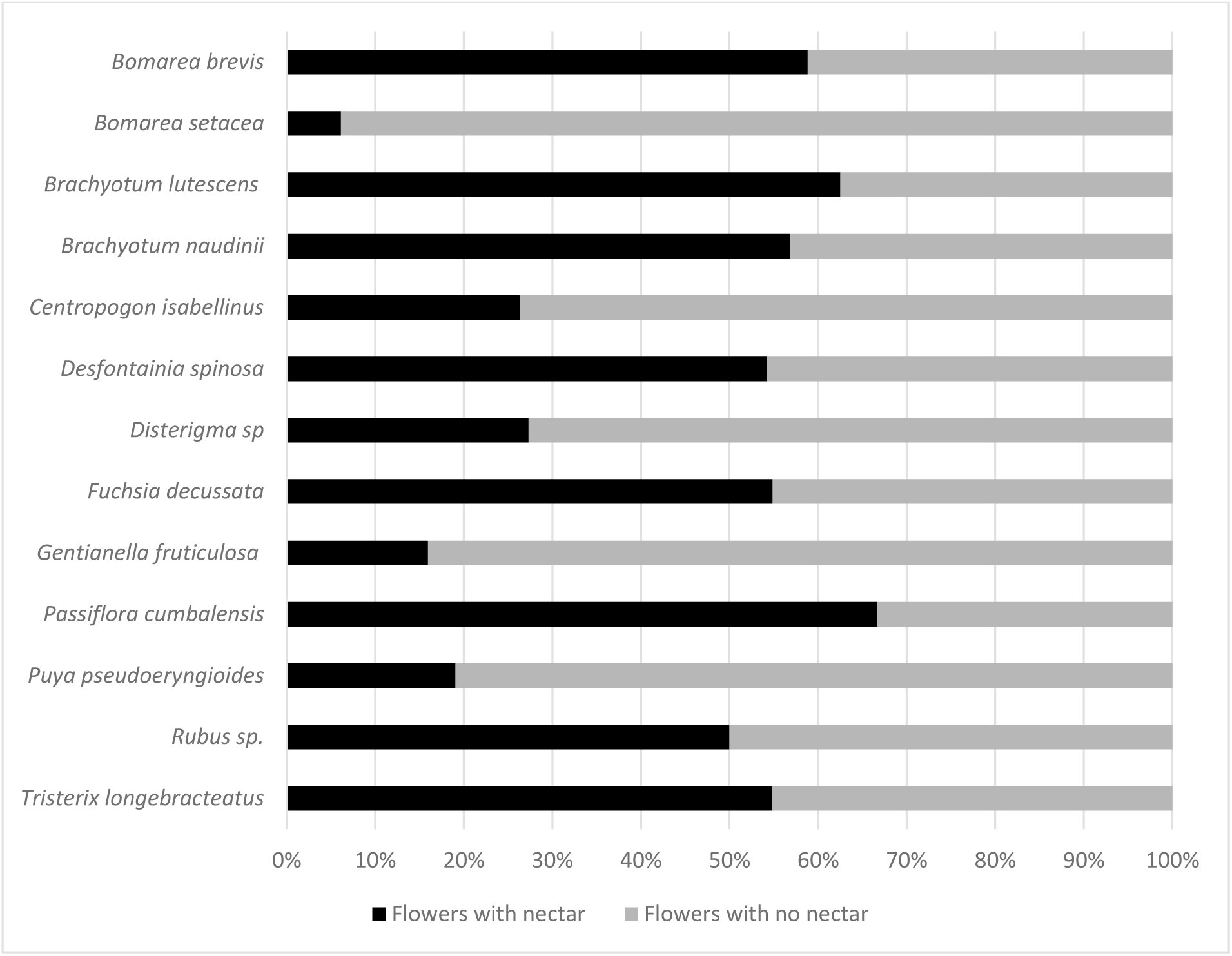
Nectar availability in flowers of plants most visited by birds in the elfin forest.

## Discussion

Plant species of elfin forests can be separated based on nectar and morphological traits (Figure 1). Half or more of the bird visits to flowers for the Metaltail, the Sapphirewing, and the Flowerpiercer were focused on plants with higher scores along the first PCA axis in the dry season (Figure 2). I expected that plants with higher energy rewards, as indicated by nectar volume and sugar concentration, would be more attractive to birds. However, this expectation held only for the Metaltail in the dry season. I found no significant association between visitation and scores along the first PCA axis, which was defined mainly by nectar rewards, for either Sapphirewings or flowerpiercers (Table 4). As the first PCA axis only captured 43-44% of the total variation, there may be other factors that are needed to explain bird visits as a function of floral characters. For bird visitors of *Rhododendron* flowers in the Himalayas, long corollas and high nectar volume are the main preferences (Basnett et al. 2019).

Plants that had both large nectar rewards and larger number of flowers per plant were the shrub *Brachyotum lutescens* and the mistletoe *Tristerix longebracteatus*. Both were frequently visited by these birds (Gonzalez and Loiselle 2016) and other nectarivorous birds in similar ecosystems, such as the elfin forest in the Colombian paramo (Gutierrez and Rojas 2001). The PCA separated plants that were more insect-pollinated than bird-pollinated; the former plants have low nectar reward and short corollas. For these plants (e.g. *Disterigma spp., Gentianella fruticulosa, Bomarea spp., Rubus*, Figure 2), the Metaltail was the more important bird visitor among those studied here (Gonzalez and Loiselle 2016).

Usually, small hummingbirds, such as the Metaltail, are generalists in terms of flower visitation while large hummingbirds like the Sapphirewing are specialists (Dalsgaard et al. 2009). The different plant species that the Metaltail and the Sapphirewing used as resources are in part similar to two of the groups of plants and hummingbirds identified by Gutierrez et al. (2004) in an elfin forest of Colombia. Small, short billed-hummingbirds tend to visit plants with a low nectar reward while large hummingbirds visit plants that have long-corolla flowers. Metaltails showed significant associations with flower characteristics along both PCA axes, which largely reflect floral rewards and flower size. Although the abundance and phenology of flowering plants varied between seasons (Fig. 3), the plant trait ordination was almost identical in both seasons (Sup. Table 1).

The Sapphirewing visited *Tristerix longebracteatus* as its primary floral resource in both dry and wet season. Other plants which had higher nectar volume and sugar content (e.g. *Puya pseudoeryngioides*, Table 2) were not visited by this bird. This result suggests that Sapphirewings might have been selecting certain plant species (e.g., *Tristerix*) rather than general plant characteristics (e.g., high energy rewards). In an elfin forest of Colombia, Sapphirewings visited primarily one *Puya* species, and such visits may be associated with plant phenology (i.e., what plants are available when birds are present) (Gutierrez et al. 2004).

The Flowerpiercer, like the Metaltail, was a generalist but tended to visit plants with high nectar reward and high number of flowers per plant, such as *Brachyotum lutescens* and *Tristerix longebracteatus.* Other species of *Diglossa* also are known to visit *Brachyotum* (Stiles et al. 1992) or *Tristerix longebracteatus* (Graves 1992). The peculiar foraging behavior of the Flowerpiercer, searching for the flowers that are on a different spatial level than flowers commonly used by hummingbirds (Feinsinger and Cowell 1978), might explain coexistence with the other two hummingbird species that use the same nectar resources. The different patterns of visitation to plants by these three bird species between seasons may be related to change in floral preferences over time (Fagua and Gonzalez 2007).

The fact that no statistical associations were detected between bird visitation and plant traits for the Metaltail in the wet season, the Sapphirewing in the dry season and the Flowerpiercer in both seasons (Table 4), suggest that other factors beyond floral traits may be needed to explain patterns of floral visitation by birds. Future studies should examine in greater detail specific preferences of bird species for plant species using controlled experiments (Maglianesi et al. 2015, Fenster et al. 2015).

## ACKNOWLEDGEMENTS

I thank Dr. Bette Loiselle for her guidance in this research. I also thank the field assistants that helped in taking several measurements of plant traits and bird visits to the plants: mainly from the Universidad de Huanuco. Funding for fieldwork in Peru came from the Premio Nacional para la Investigacion Ambiental of the Ministerio del Ambiente of Peru, Optics for the Tropics, Royal Society for Bird Protection, Idea Wild and University of Florida’s Tropical Conservation and Development Program field research fund. Research Grant. Thanks to FINCyT in Peru and WWF for funding my PhD studies.

---

**Supplementary Table 1.** Principal component analysis for the plant traits that are visited by the most connected birds in the bird-flowering plant visitation network.

|  | Dry Season |  | Wet Season |  |
| --- | --- | --- | --- | --- |
|  | Axis 1 | Axis 2 | Axis 1 | Axis 2 |
| Eigenvalue | 4.328 | 2.296 | 4.445 | 2.168 |
| % of var. | 43.276 | 22.959 | 44.445 | 21.678 |
| Cumulative % of var. | 43.276 | 66.235 | 44.445 | 66.123 |
| PC1 | correlation | p value | correlation | p value |
| Max nectar | 0.941 | * | 0.940 | * |
| Nectar per flower | 0.940 | * | 0.955 | * |
| Milligrams sugar per flower | 0.886 | * | 0.906 | * |
| Maximum nectar concentration | 0.823 | * | 0.774 | * |
| Flowers per individual | 0.623 | 0.02 | 0.676 | 0.01 |
| Height of flowers | 0.508 | 0.07 | 0.525 | 0.11 |
| Corolla length | 0.446 | 0.12 | 0.456 | 0.11 |
| C.V. Milligrams sugar per flower | 0.379 | 0.2 | 0.459 | 0.11 |
| C.V. Microliters nectar per flower | 0.152 | 0.6 | 0.126 | 0.68 |
| Corolla width | -0.287 | 0.3 | -0.246 | 0.41 |
| PC2 |  |  |  |  |
| Corolla length | 0.849 | * | 0.844 | * |
| Height | 0.661 | 0.01 | 0.629 | 0.02 |
| Flowers per individual | -0.619 | 0.02 | -0.632 | 0.02 |
| Corolla width | 0.524 | 0.06 | 0.553 | 0.04 |
| Maximum nectar concentration | 0.291 | 0.33 | 0.337 | 0.25 |
| Milligrams sugar per flower | 0.201 | 0.50 | 0.095 | 0.75 |
| Maximum nectar concentration | -0.164 | 0.59 | -0.163 | 0.59 |
| C.V. Microliters nectar per flower | -0.222 | 0.46 | -0.251 | 0.40 |
| Nectar per flower | -0.246 | 0.41 | -0.169 | 0.58 |
| C.V. Milligrams sugar per flower | -0.465 | 0.10 | -0.334 | 0.26 |

\*p value < 0.01

